# The Inferior Frontal Sulcus, not the Anterior Insula, acts as a late-stage attentional gate for the fast rejection of salient but irrelevant sensory inputs during cognitive tasks

**DOI:** 10.64898/2025.12.15.694314

**Authors:** Benoit Chatard, Maryne Dupin, Mathilde Petton, Benjamin Bontemps, Lorella Minotti, Philippe Kahane, Sylvain Rheims, Jean-Philippe Lachaux

**Affiliations:** Université Claude Bernard Lyon 1, CNRS, INSERM, Centre de Recherche en Neurosciences de Lyon CRNL U1028; UMR5292, EDUWELL, F-69500, Bron, France; Univ. Grenoble Alpes, Inserm, U1216, CHU Grenoble Alpes, Grenoble Institut Neurosciences, 38000 Grenoble, France; Neurology Department, University Hospital of Grenoble, Grenoble, France; Lyon Neuroscience Research Center (CRNL, INSERM U1028/CNRS UMR 5292, Lyon 1 University), Lyon, France; Department of Functional Neurology and Epileptology, Hospices Civils de Lyon and Lyon 1 University, Lyon, France

## Abstract

The ability to prioritize behaviorally relevant stimuli is essential for efficient cognition, particularly in distracting environments. The anterior insula (AI), a key node of the salience network, has been proposed to support this selection process by detecting salient events. A similar function, however, has also been attributed to the inferior frontal sulcus (IFS), located at the intersection of the ventral and dorsal attention networks. Here, we directly compared the respective contributions of these regions to attentional gatekeeping—the identification and rejection of attention-grabbing yet task-irrelevant “false positives” to preserve cognitive resources. We recorded intracranial EEG from 63 patients performing an attentive reading task designed to dissociate salience from task relevance. High-Frequency Activity (HFA) responses between 50 Hz and 150 Hz revealed that the IFS was the only region—across an extensive sampling of the frontal and insular cortices—to exhibit consistent, subsecond response dynamics compatible with the filtering of false positives. In contrast, the AI did not participate in the analysis or rejection of distractors and showed only sparse, delayed responses to targets, suggesting a role in coordinating or energizing downstream cognitive or motor processes rather than in salience detection per se. These findings indicate that the IFS—rather than the AI—is the primary late-stage node for attentional gatekeeping during cognitive tasks performed under distracting conditions.

## Introduction

Because the human brain has limited capacity to process sensory inputs, attentional selection is continuously required to prioritize stimuli with the highest behavioral relevance (***Duncan (1980***), ***Fiebelkorn and Kastner (2019***)). To be effective, this selection must minimize both false negatives—failing to detect important stimuli—and false positives—allocating cognitive resources to irrelevant ones (***Kastner and Ungerleider, 2000***). A familiar example of false-positive rejection is ignoring an email alert that momentarily captures our gaze while drafting a manuscript. This ability to suppress responses to attention-grabbing but irrelevant stimuli relies on a “gate-keeping” mechanism that evaluates stimulus relevance in the context of the current task set (***Sakai, 2008***).

During a task, sensory systems can be fine-tuned to respond maximally to features shared by goal-relevant stimuli—as encoded in the attentional set or attentional templates—to limit the likelihood of both false negatives and false positives (***Theeuwes (1994***); ***Dosher and Lu (2000***), ***Maunsell and Treue (2006***), ***Treue and Martinez Trujillo (1999***)) According to one recent theoretical proposal—the Unified Diachronic Account of Attentional Selectivity (UDAS)—such tuning increases the probability that stimuli with desired features cross a critical threshold and trigger an “attentional episode” characterized by access to working memory and response-related processes (***Zivony and Eimer, 2022***).

Outside of specific task contexts, behavioral relevance is determined by a loose hierarchy of concurrent goals, ranging from maintaining physical comfort to preserving social status (***Peelen and Kastner, 2014***). In these situations, the transient sensory tuning proposed by the UDAS cannot fully account for attentional selection across the wide variety of potentially relevant stimuli, indicating that additional mechanisms must be involved. The salience network (SN), anchored in the anterior insula (AI) and anterior cingulate cortex (ACC), has been proposed to detect stimuli with high homeostatic relevance (***Seeley, 2019***), while the ventral attention network (VAN)—a right-lateralized frontoparietal system that also includes the AI—has been associated with the reorientation of attention toward behaviorally significant events (***Corbetta and Shulman, 2002***).

Because the VAN and SN are also active during tasks with well-defined attentional sets, they are expected to interact closely during the selection of task-relevant stimuli, with precise timing relative to attentional episodes. Yet, the degree of functional dissociation between these networks, and their respective dynamics in response to incoming stimuli, remains poorly understood. Corbetta and Shulman proposed that many task-irrelevant stimuli may not reach the VAN due to an early gating mechanism that discards potential false negatives based on low-level features (***Corbetta and Shulman, 2002***), a model compatible with the thresholding framework of the UDAS. In this view, the VAN (and possibly the SN) would process only those stimuli that have already triggered an attentional episode, applying a more elaborate match with the attentional set to provide a secondary gating mechanism specifically aimed at avoiding false positives.

One approach to anatomically localize such late-stage attentional gating is to record whole-brain activity during paradigms that present attention-capturing stimuli capable of passing early filters, with some requiring extensive cognitive processing (targets) and others to be ignored through a late filtering stage (distractors). Regions involved in false-positive rejection should respond similarly to all stimuli prior to the “divergence latency,” the moment when target- and distractor-evoked activity begins to differ due to target-specific processes.

Using this approach, prior work using the high spatiotemporal resolution of intracerebral EEG (iEEG) identified a region within the VAN—near the inferior frontal sulcus (IFS)—whose response properties are consistent with a late gate-keeping mechanism (***Perrone-Bertolotti and others, 2020***). The present study builds on this framework to examine the respective contributions and temporal dynamics of the VAN and SN during false-positive rejection, with particular attention to the anterior insula (AI)—given its involvement in both networks—and the IFS. Based on extensive iEEG sampling of the frontal lobe in 63 patients, we show that only the IFS, not the AI, exhibits response dynamics consistent with a role in attentional selection.

## Results

Following the original design of the Attentive Reading Task (***Nobre et al., 1998***) (***Figure 1***), we first verified that all patients had correctly read the target story (grey words) by asking comprehension questions immediately after the task. All participants answered accurately, and none reported noticing that distractor words occasionally formed incoherent sentences—a feature that would have been salient to anyone attending to the non-target stream. When explicitly asked whether they had read both stories, all participants stated that doing so would have been impossible and recalled being instructed to focus exclusively on the target story. Taken together, these behavioral confirmations, along with the electrophysiological data described below, indicate that the task was performed as intended.

**Figure 1.**
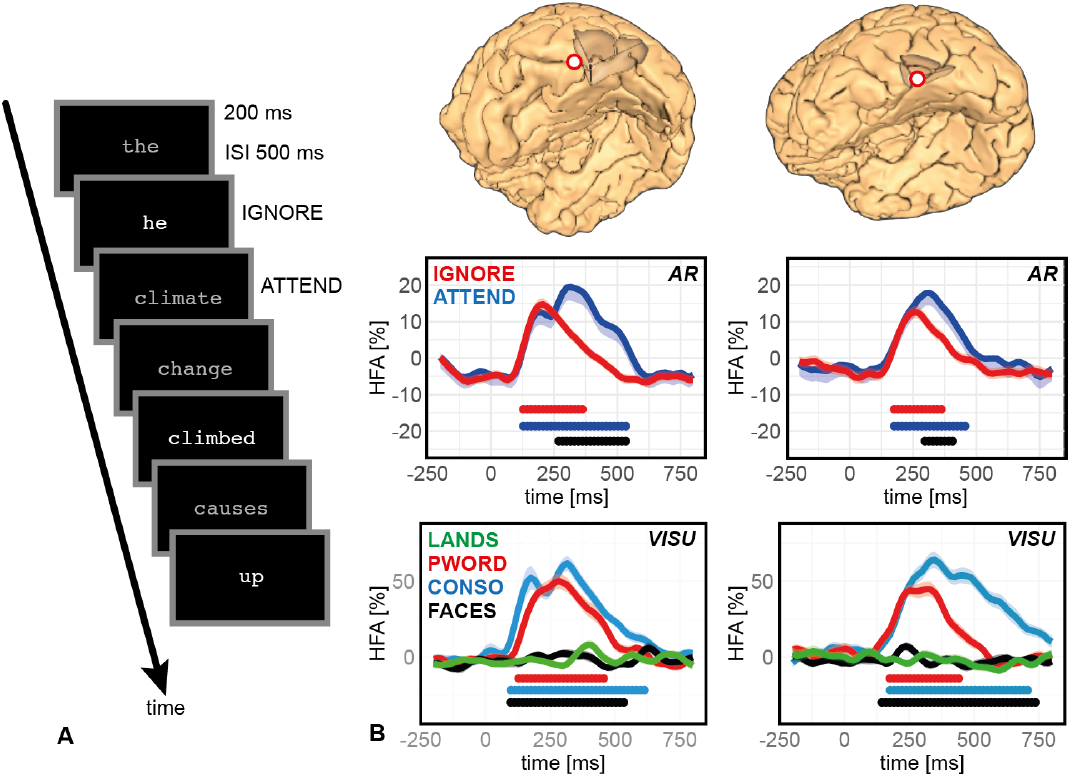
Experimental design of the Attentive Reading Task and estimation of Divergence Latency. Left Panel: Attentive Reading (AR) task : participants read two intermixed stories presented word by word (200 ms per word, 700 ms interstimulus interval) and attended to the gray (low contrast) story while ignoring the white (high contrast) distractors. Each block comprised 400 words, with a pseudo-randomized color distribution. Right Panel: Illustrations of two sites within the Visual Word Form Area (VWFA) mapped onto the individual anatomy of two patients (red dots), depicting high-frequency activity (HFA, between 50 Hz and 150 Hz) during the AR task and the visual oddball task (VISU). HFA is expressed as a percentage deviation from the mean HFA observed during the entire experiment for each specific iEEG site. Colored horizontal bars indicate episodes of significant HFA increase relative to the pre-stimulus baseline [-200:0 ms], sustained for over 100 ms (stimulus onset at 0 ms; Wilcoxon, p < 0.001, uncorrected). The black horizontal bar indicates significant differences in HFA between experimental conditions lasting more than 100 ms (Kruskal-Wallis, p < 0.001, uncorrected). VWFA sites exhibited a selective HFA increase for pseudowords and consonant strings (PWORD and CONSO), with no response to face and landscape stimuli (FACES and LANDS). The divergence latency was established at 250 ms, corresponding to the differential response to attended words (ATTEND) versus ignored words (IGNORE) in the AR task.

### Finding the divergence latency

Our approach required a precise estimate of the latency at which responses to distractor and target words began to diverge, used here as a proxy for the moment when the decision to trigger a selective response to targets is made. This divergence latency was assessed using intracerebral recordings from the visual word form area (VWFA), localized in the left basal temporal lobe based on its responses to word and non-word stimuli in the visual oddball task (***Figure 1***). The MNI coordinates of the two sites ([-45;-56;-4] and [-46;-57;-15]) were consistent with previous descriptions of the VWFA (***Chen et al., 2019***).

During the Attentive Reading task, the VWFA exhibited an early, non-selective peak at approximately 200 ms in response to both target and distractor words, followed by a sustained targetselective component lasting until 500–600 ms. The onset of this target-specific activity was estimated at 250 ms, based on a statistical comparison between conditions (ATTEND vs. IGNORE; Kruskal–Wallis test, p < 0.001, uncorrected). This estimate was further supported by recordings from additional sites showing target-selective responses, including a region associated with covert speech in the precentral gyrus (data not shown).

### Early Undifferentiated Responses in the Frontal lobe

The next step was to identify sites that responded to both targets and distractors before the divergence latency of 250 ms. To isolate regions engaged in early, non-selective attentional processing, we examined responses to both stimulus types within this time window. Analyses were restricted to frontal and insular cortices anterior to the central sulcus, encompassing 4,236 iEEG sites (2,263 in the left hemisphere; 1,973 in the right; ***Figure 2***). Sites showing a significant increase in high-frequency activity (HFA) for both targets and distractors prior to 250 ms were identified using a Wilcoxon test relative to the pre-stimulus base-line (200 to 0 ms; P < 0.001, uncorrected). Uncorrected thresholds were used to maximize sensitivity and avoid excluding regions potentially involved in gate-keeping mechanisms. Thirty-six sites met our criteria for an “early undifferentiated response” (EUR), defined by: (i) a response to both stimulus types, (ii) onset before 250 ms, and (iii) no significant difference between conditions prior to this latency (Kruskal–Wallis test, P > 0.001, uncorrected). These EUR sites were distributed across 17 patients (29 in the left hemisphere; 7 in the right). To avoid redundancy arising from adjacent responsive contacts along the same electrode, we applied a single-strongest-response (SSR) criterion, retaining only the most responsive site per electrode per patient. This yielded 22 unique EURs (19 left hemisphere; 3 right; Figure B).

**Figure 2.**
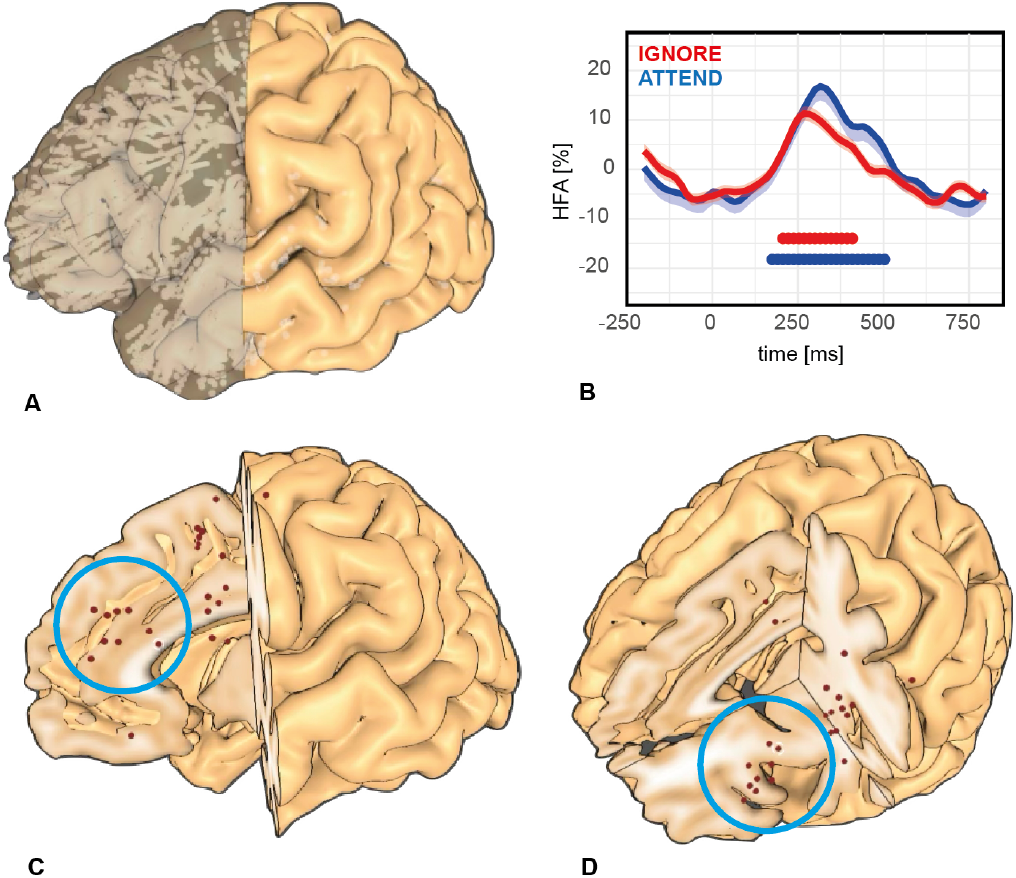
Global iEEG sampling and early undifferentiated responses (EURs). A: Distribution of all frontal iEEG recording sites (white dots) across 63 patients. B: Example of an Early Undifferentiated Response (EUR) from a single site, showing a significant increase in high-frequency activity (HFA) relative to the pre-stimulus baseline, beginning before 250 ms for both attended (ATTEND) and ignored (IGNORE) words in the Attentive Reading task. Colored horizontal bars indicate periods of significant HFA increase relative to baseline (200 to 0 ms) lasting more than 100 ms (Wilcoxon test, p < 0.001, uncorrected; stimulus onset at 0 ms). HFA is expressed as a percentage deviation from the session-wide mean. C-D: All sites showing an EUR (Single Strongest Response per electrode) in the frontal or insular cortex, projected onto a 3D MNI brain. The blue circle marks EURs located in the inferior frontal sulcus.

### Early Undifferentiated Responses in the frontal eye field

Most early undifferentiated responses (EURs) were located in the precentral gyrus and sulcus, particularly within motor regions involved in the preparation and execution of eye, hand, and laryngeal movements. Given the substantial inter-individual variability in the motorotopic organization of these areas, we used two additional tasks—a visual search task (SEARCH) and a phonological task (***Mainy et al., 2008***)— (***Figure 1***) to differentiate sites involved in attentional processing (i.e., the frontal eye field, FEF) from those related to speech-related motor activity. ***Figure 3***) shows selective anatomical activation of the FEF during visual search.

**Figure 3.**
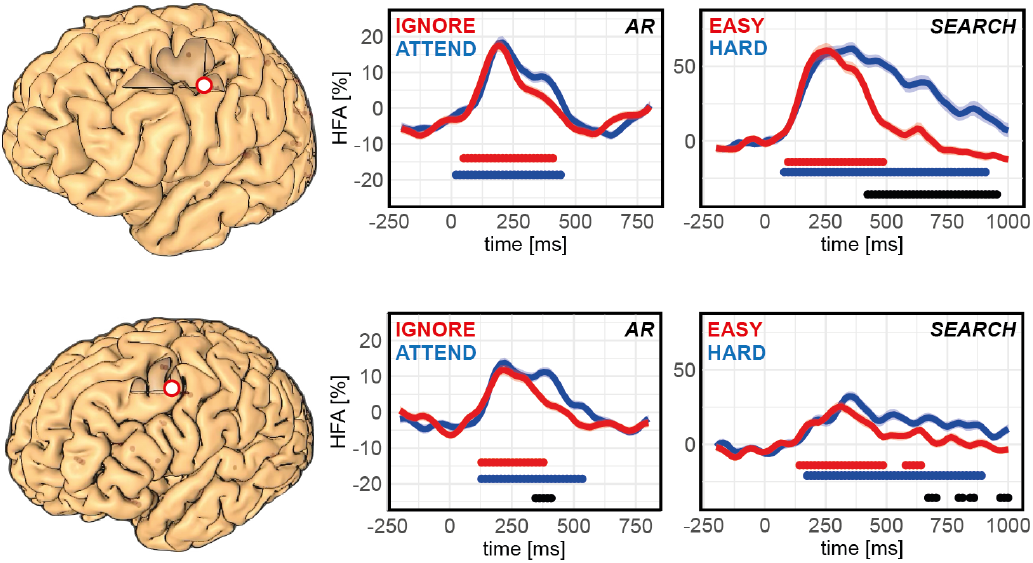
Early responses to attended and ignored words in the frontal eye field (FEF). Two representative FEF sites are shown on individual 3D brain reconstructions (red dots). Both exhibited an Early Undifferentiated Response (EUR) during the Attentive Reading task, with onsets earlier than 250 ms (middle panels). Colored horizontal bars indicate periods of significant high-frequency activity (HFA) increase relative to the pre-stimulus baseline (-200 to 0 ms), lasting more than 100 ms (Wilcoxon test, p < 0.001, uncorrected; stimulus onset at 0 ms). HFA is expressed as a percentage deviation from the mean HFA across the recording session. Localization within the FEF was confirmed using a visual search task, which evoked sustained activity in both sites, with a longer HFA increase in the difficult search condition (target and distractors of the same color) compared to the easy condition (color pop-out) (right panels).

To focus on attentional rather than motor responses, we excluded sites associated with hand or laryngeal movements and concentrated on the FEF (***Derrfuss et al., 2004***), a key node of the dorsal attention network implicated in the rapid reorientation of attention to salient stimuli. We identified three sites (based on the single strongest response, SSR) whose anatomical and functional profiles were consistent with the FEF (including a sustained activity throughout the visual scan of the stimuli). All three exhibited early, non-selective response peaks to both targets and distractors at approximately 200 ms—well before the divergence latency. Representative examples are shown in ***Figure 3***.

### An accumulation of Early Undifferentiated Responses (EUR) near the inferior frontal sulcus

In addition to the precentral gyrus and sulcus, we identified 19 EUR sites outside motor regions (11 in the left hemisphere; after applying the SSR procedure: 12 sites across 12 patients—8 in the left hemisphere, 4 in the right). Notably, these EURs clustered in or near the inferior frontal sulcus (IFS), comprising 5 of the 8 left-hemisphere sites and all 4 right-hemisphere sites (SSR). ***Figure 4*** illustrates six representative sites projected onto individual brain surfaces : despite extensive sampling across the frontal lobe, these responses were strikingly localized (see Supplementary Figure 1 for additional examples).

**Figure 4.**
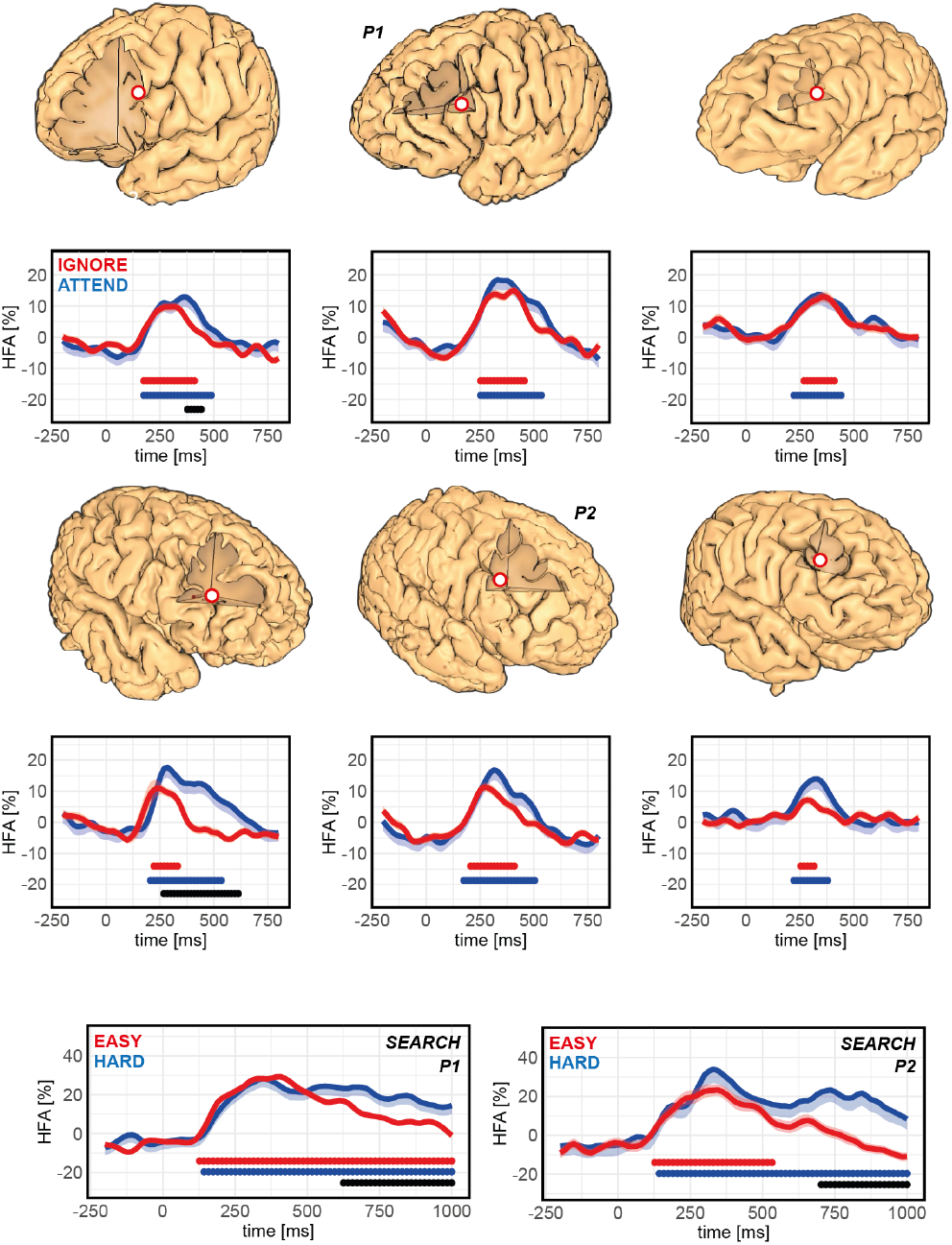
Early Responses to Attended and Ignored Words in the Inferior Frontal Sulcus. This figure illustrates the response dynamics for six representative iEEG sites located in the Inferior Frontal Sulcus (IFS), displayed on individual 3D brain models (red dots). All sites exhibit an Early Undifferentiated Response during the Attentive Reading task. Colored horizontal bars indicate episodes of significant HFA increase relative to the pre-stimulus baseline [-200:0 ms], sustained for over 100 ms (stimulus onset at 0 ms; Wilcoxon, p < 0.001, uncorrected). HFA is expressed as a percentage deviation from the mean HFA recorded throughout the experiment. Bottom row : HFA response in the IFS during the visual search task for sites P1 and P2, showing a clear and early activity increase in this non-verbal paradigm.

Although observed in only nine patients, the EURs represented the majority of recorded sites within this well-defined anatomical region. A detailed analysis of all sites within an MNI-defined bounding box surrounding the nine IFS EURs (X > 35 or X < –35; 15 < Y < 35; 10 < Z < 25) revealed only six additional sites without EURs (Supplementary Figure 2), indicating that most sites sampled in this region (9/15) exhibited an early, undifferentiated response.

### Early undifferentiated responses in the preSMA

In addition to the inferior frontal sulcus (IFS), the only frontal region exhibiting early undifferentiated responses (EURs) in more than one patient was the Supplementary/PreSupplementary Motor Area (SMA/preSMA), with two sites identified in two left-hemisphere patients. Given the proximity of this region to the mid-ACC—a core node of the salience network—we carefully examined the anatomical locations of these two sites. Both were confirmed to lie above the cingulate sulcus, at the boundary between the SMA and preSMA (***Figure 5***). A systematic inspection of all 189 recording sites within the anterior medial frontal wall (97 in the left hemisphere) revealed no EURs within the ACC proper. One additional EUR site was identified in the lateral orbitofrontal cortex of a single patient.

**Figure 5.**
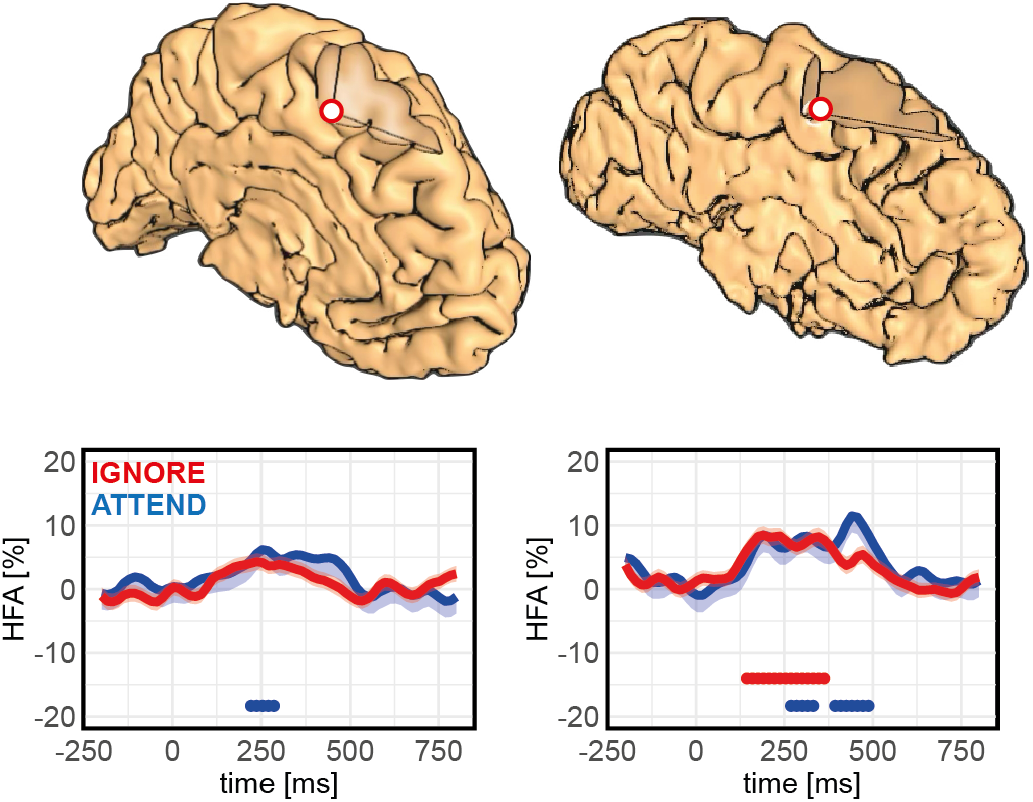
Early responses to attended and ignored words in the SMA/pre-SMA. High-frequency activity (HFA) responses from two sites located at the boundary between the Supplementary Motor Area (SMA) and pre-SMA are shown on the pa-tient’s individual anatomy (red dot). The sites exhibit a clear HFA peak before 250 ms, with comparable amplitudes for attended (ATTEND) and ignored (IGNORE) words. Colored horizontal bars indicate periods of significant HFA increase relative to the pre-stimulus baseline (200 to 0 ms), lasting more than 100 ms (Wilcoxon test, p < 0.001, uncorrected; stimulus onset at 0 ms). HFA is expressed as a percen-tage deviation from the session-wide mean. Note : In the left panel, the IGNORE condition shows a significant HFA increase before 250 ms, but it does not appear as a colored bar because its duration [75 ms] did not meet the 100 ms threshold.

### Absence of early undifferentiated responses in the anterior insula

Given that distractors in the Attentive Reading task were intentionally designed to be highly salient and attention-capturing, we hypothesized that regions implicated in salience detection—particularly the anterior insula—would exhibit early undifferentiated responses (EURs). To test this, we systematically applied the EUR detection procedure to all anterior insular recording sites, distinguishing dorsal (dAI) from ventral (vAI) subdivisions based on visual inspection of each patient’s 3D anatomical insula reconstruction using HiBoP (***Vecchio et al., 2024***). Defining the anterior insula as the region encompassing the anterior and middle short gyri, we identified 92 sites in the dAI and 33 in the vAI. None met the criteria for an EUR. More broadly, only four anterior insular sites showed any significant task-related response, and three of these were selective for target words (***Figure 6***). All responsive sites were located in the dorsal portion of the anterior short gyrus.

**Table 1.** Number of iEEG sites showing a significant HFA increase relative to the pre-stimulus baseline (−200 to 0 ms; Wilcoxon, *p <* 0.001, uncorrected) in response to attended words (ATTEND), distractor words (IGNORE), or both, relative to the total number of sites recorded in each insula subregion (v/d: ventral/dorsal; A/M/P: Anterior/Middle/Posterior; S/L: Short/Long; G: Gyrus).

| Region | vASG | dASG | vMSG | dMSG | vPSG | dPSG | vALG | dALG | vPLG | dPLG |
| --- | --- | --- | --- | --- | --- | --- | --- | --- | --- | --- |
| ATTEND | 0 | 3 | 0 | 0 | 0 | 0 | 0 | 0 | 0 | 0 |
| IGNORE | 0 | 1 | 0 | 0 | 0 | 0 | 1 | 1 | 0 | 0 |
| BOTH | 0 | 0 | 0 | 0 | 0 | 0 | 0 | 0 | 0 | 0 |
| # SITES | 13 | 46 | 20 | 46 | 22 | 35 | 26 | 56 | 33 | 27 |

**Figure 6.**
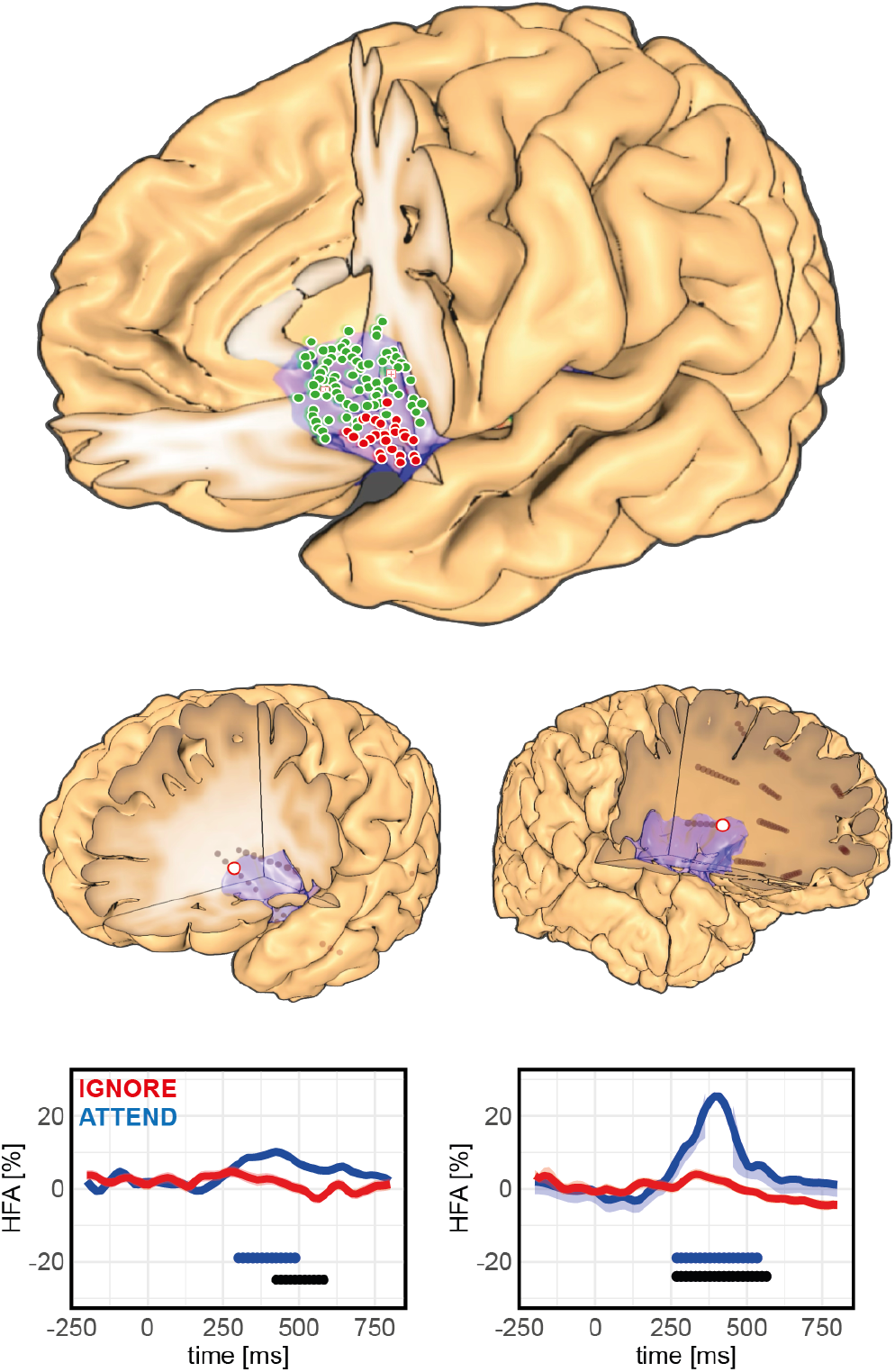
Late responses selective to attended words in the anterior insula. Top: Distribution of all iEEG recording sites within the dorsal (green dots) and ventral (red dots) anterior insula across 63 patients. Bottom: High-frequency activity (HFA) responses from two sites located in the dorsal anterior insula (anterior short gyrus), shown on individual 3D brain reconstructions (red dots). Colored horizontal bars indicate periods of significant HFA increase relative to the pre-stimulus baseline (-200 to 0 ms), lasting more than 100 ms (Wilcoxon test, p < 0.001, uncorrected; stimulus onset at 0 ms). Black horizontal bars denote significant differences between experimental conditions lasting more than 100 ms (Kruskal-Wallis test, p < 0.001, uncorrected). HFA is expressed as a percentage deviation from the mean activity across the recording session.

### Comparing response latencies within individuals

Intracerebral EEG (iEEG) recordings occasionally offer the unique opportunity to measure neu-ral responses simultaneously across multiple regions of interest within the same individual, allowing direct comparisons of response latencies. Supplementary Figure 3 illustrates such a case, with concurrent recordings from the frontal eye field (FEF), inferior frontal sulcus (IFS), and visual word form area (VWFA) in a single patient. The FEF exhibited early responses to both targets and distractors, with onsets preceding 200 ms. This activity temporally preceded the undifferentiated response observed in the IFS, which itself occurred before the divergence latency identified in the VWFA. These within-subject comparisons support a temporal cascade of activation consistent with a hierarchical attentional selection process.

## Discussion

Even during demanding tasks, attention is often momentarily diverted by salient distractors. Although such attentional capture is typically brief (***Huang and Yeh, 2011***), it is essential that cognitive resources be rapidly disengaged to remain available for more relevant stimuli. Our aim was to investigate how the brain handles these “false positives,” focusing on two networks frequently implicated in the detection of behaviorally significant information: the ventral attention network (VAN) and the salience network (SN), and within these networks, the inferior frontal sulcus (IFS) and the anterior insula (AI).

We used a reading task in which both target and distractor words were designed to capture attention. Successful performance required selectively engaging high-level processing for targets while suppressing processing of distractors—a function we refer to as “gate-keeping.” By estimating the latency of the target-specific response in the visual word form area (VWFA), we inferred that this selection occurs before 250 ms (the “divergence latency”). Across the prefrontal and insular cortices, only the inferior frontal sulcus (IFS) consistently exhibited response dynamics compatible with this selective gate-keeping.

### Revisiting the Role of the Anterior Insula in Salience Detection

In contrast to the IFS, we found no evidence that the anterior insula responds to both targets and distractors before the divergence latency (250 ms), suggesting that it does not contribute to the late-stage filtering of false positives. Although we cannot entirely exclude the existence of a small, unsampled region that might play such a role, this seems unlikely given the limited size of the anterior insula—only a few cubic centimeters—and the extensive coverage provided by our dataset (126 recording sites, each sampling a radius of approximately 3 mm; (***Lachaux et al., 2003***)). A similar sampling argument was used by Woolnough et al. (***Woolnough et al., 2019***) to rule out a pre-articulatory role of the insula in language production.

Our findings also challenge the idea that the anterior insula is centrally involved in preventing false negatives, at least under our experimental conditions. If the AI ensured that all potentially important stimuli were processed—a key function attributed to the salience network—it should have responded to both targets and distractors in our reading task. Instead, most AI sites showed no detectable response. These results suggest that the AI does not contribute to the detection of behaviorally relevant stimuli when attentional sets are well defined. Nevertheless, we cannot formally rule out a role in detecting salient events outside of a structured task context, in line with broader accounts of salience network function (***Seeley and others, 2007***). One possibility, for instance, is that salience detection in the AI may be suppressed for stimuli that are repeatedly presented in a fixed location.

In the dorsal AI, however, this interpretation is more difficult to reconcile with our observation that a subset of sites responded selectively to target stimuli at relatively late latencies, after the IFS. This pattern aligns more closely with recent proposals that the dorsal AI contributes to the coordination of task-related responses within the so-called Action Mode Network (***Dosenbach et al., 2025***), after false positives have been filtered out by a gate-keeping mechanism. Even so, this interpretation must be tempered by the fact that most dorsal AI sites showed no such response. A plausible explanation for this counterintuitive result is that, in skilled readers, word processing is highly automatized, and recruitment of the dorsal AI becomes necessary only when cognitive control must override default responses. This would imply that the dAI is engaged when a specific response-related network must be activated—but not inhibited—since in our task, the inhibition of reading automatisms was required for distractors.

### A late attentional gate-keeping mechanism in the IFS, at the intersection of the VAN and DAN

Our findings provide strong support for the involvement of the inferior frontal sulcus (IFS) in detecting behaviorally relevant stimuli, likely within a network that includes other regions showing early, undifferentiated responses to both targets and distractors (such as the preSMA). Although our data cannot definitively establish that this network evaluates the behavioral relevance of stimuli that have already captured attention, this interpretation is consistent with previous evidence implicating the lower portion of the dorsolateral prefrontal cortex—including the IFS—in maintaining the attentional set, shifting attention to relevant stimulus dimensions, and discriminating between task-relevant and task-irrelevant inputs when the distinction is difficult (***Derrfuss et al. (2004***),***Milham and others (2001***),***Zysset and others (2001***)). Notably, we observed no hemispheric asymmetry in early IFS responses, and clear activations in a nonverbal visual search task in both hemispheres. This challenges earlier proposals of a functional specialization whereby the right IFS supports attentional control and the left IFS supports language processes (***Ruland et al., 2022***).

The anatomical and functional position of the IFS—at the intersection of ventral and dorsal lateral prefrontal cortex (***Ruland et al., 2022***) and at the boundary between the Ventral and Dorsal Attention Networks (VAN and DAN; (***Corbetta and Shulman, 2002***))—raises important questions about its precise contributions to these networks. The VAN is typically recruited by task-irrelevant but salient stimuli, such as the distractors used in our Attentive Reading task. Thus, the presence of robust responses to both targets and distractors around the IFS suggests that this region represents a key node of the VAN. One might object that the VAN has been predominantly described as right-lateralized, whereas some of the clearest IFS responses in our study were observed in the left hemisphere. However, a careful reading of the foundational VAN literature (***Corbetta et al. (2008***),***Corbetta and Shulman (2002***)) reveals no strong claim of complete lateralization, leaving open the possibility that the left IFS also forms part of this network.

An important remaining question is whether the IFS should also be considered part of the DAN. Corbetta and Shulman (***Corbetta and Shulman, 2002***) included a region in the posterior middle frontal gyrus, near the IFS, within both the VAN and the DAN. The IFS may therefore also contribute to the goal-directed, top-down allocation of attention attributed to the DAN.

### A gating mechanism compatible with rhythmic attentional sampling

The timing of activation in the IFS provides critical insights into its role in attentional control, particularly within the framework of the Unified Diachronic Account of Attentional Selectivity (UDAS; (***Zivony and Eimer, 2022***)). According to this model, perceptual evidence accumulates within sensory systems that are biased by the current task set. When this activity reaches a critical threshold, it triggers a transient “attentional episode,” during which selected stimuli gain access to working memory and response-related processes. A key strength of the UDAS is that it specifies expected activation latencies for several anatomical regions. In particular, the frontal eye field (FEF) is predicted to be active during the attentional episode, which is thought to occur after approximately 170 ms. In one patient, we were able to record simultaneously from the FEF, the visual word form area (VWFA), and the IFS, and the observed timings were fully consistent with the UDAS predictions: FEF activation occurred before 200 ms. The VWFA showed two distinct components—an early response aligned with FEF activity, and a later, target-specific component occurring after the IFS peak. Most strikingly, the IFS became active after the FEF peak, corresponding—within the UDAS frame-work—to the moment when attention has been captured during the attentional episode. Because the IFS peak also preceded the target-specific VWFA response, the FEF–IFS–VWFA sequence supports the interpretation that the IFS participates in the decision process that determines whether an attended stimulus should receive further processing (***Cole and Schneider, 2007***). Thus, the IFS appears to be an integral component of the attentional selection system, operating on stimuli that have already captured attention. Interestingly, the timing of IFS activity is also compatible with another influential account of attention—the rhythmic model (***Fiebelkorn and Kastner, 2019***)—which proposes that attention samples the environment in cycles as short as 250 ms. A mechanism evaluating the behavioral relevance of each attended item within a 250–300 ms window, involving the IFS, fits naturally within such rhythmic sampling dynamics.

## Conclusion

We conclude that although parts of the anterior insula may participate in detecting salient stimuli—particularly those that are unpredictable or outside the current attentional focus—its dorsal portion is more likely involved in orchestrating or energizing the response to such stimuli, at least when that response requires executive control. The prevention of false positives, at least in the context of a well-defined task, appears instead to rely on the inferior frontal sulcus (IFS), located at the interface between the VAN and the DAN in both hemispheres. This region responds to stimuli that have already captured attention and seems to contribute to the decision of whether such stimuli should undergo further processing. Finally, we acknowledge that our conclusions are based solely on analyses of high-frequency activity (50–150 Hz), which may be viewed as a limitation. However, this choice was motivated by the fact that HFA serves as a reliable proxy for population-level spiking activity (***Buzsáki et al., 2012***). Thus, any cortical structure transmitting information to response-related areas about the relevance of incoming stimuli would be expected to generate a detectable increase in HFA.

## Materials and Methods

### Participants and Recordings

Intracranial electroencephalography (iEEG) recordings were obtained from 63 neurosurgical patients with pharmacoresistant epilepsy at the Epilepsy Departments of the Grenoble University Hospital and the Lyon Neurological Hospital. All participants were stereotactically implanted with multilead depth electrodes as part of their presurgical evaluation. Implantation sites were determined solely on clinical grounds following standard medical procedures. Electrode contacts located within seizure onset zones or exhibiting significant interictal spiking were excluded from all analyses. All participants provided written informed consent, and the study protocol was approved by the Institutional Review Board and the National French Science Ethical Committee (CPPRB; CPP Sud-Est V 09-CHU-12). All participants had normal or corrected-to-normal vision and were native French speakers. Between eleven and fifteen semirigid multilead electrodes were implanted in each patient. These stereotactic EEG electrodes (DIXI Medical) have a diameter of 0.8 mm and consist of 10 to 15 contacts, each 2 mm in length and spaced 1.5 mm apart, depending on the target structure. Electrode contacts were first identified on the individual implantation scheme and then anatomically localized using Talairach and Tournoux’s proportional atlas (***Talairach and Tournoux, 1993***). Co-registration between post-implantation CT and pre-implantation 3D MRI scans (VOXIM R, IVS Solutions), or between post- and pre-implantation MRIs (using IntrAnat; (***Deman and others, 2018***)), enabled precise mapping of electrode contacts onto each patient’s anatomy. Electrode visualization on individual brains and on the MNI template was performed using HiBOP (***Vecchio et al., 2024***).

Intracranial EEG signals were recorded using a video-iEEG monitoring system (Micromed), allowing simultaneous acquisition from up to 128 depth electrode contacts. Signals were band-pass-filtered online between 0.1 and 200 Hz and sampled at either 512 or 1024 Hz. During acquisition, recordings were referenced to a contact located in the white matter; for all subsequent analyses, signals were re-referenced to their immediate neighbors using a bipolar montage to improve spatial specificity and reduce common-mode noise (***Mercier and others (2022***);***Ossandon and others (2012***)).

### Experimental tasks and stimuli

#### Attentive Reading Task (AR)

This task was adapted from (***Nobre et al., 1998***). In each experimental block, participants viewed two intermixed stories presented word by word at a rapid pace (200 ms per word, with a 700 ms interstimulus interval; ***Figure 1***). Against a black background, one story appeared in grey and the other in white. Consecutive words of the same color formed a coherent, simple story in French. Participants were instructed to read the grey (low-contrast) story and report it at the end of each block, while ignoring the white (high-contrast) words. The color scheme was chosen to reduce the efficiency of early filtering mechanisms based solely on physical features, as both stories appeared in the same spatial location and distractors were displayed with higher contrast. Each block contained 400 words: 200 grey words (task-relevant ATTEND stimuli) and 200 white words (task-irrelevant IGNORE stimuli). Color assignment was pseudo-randomized to prevent predictability and to ensure that no more than three consecutive words shared the same color. At the end of the experiment, participants answered comprehension questions about the attended story, requiring specific recall of its content. Some words from the ignored story were deliberately shu?ed to create inconsistencies for participants who might have inadvertently attended to it. All words were presented foveally, and participants could not anticipate the color of the upcoming word, ensuring that every stimulus captured visual attention. While early filtering could theoretically favor targets based on physical characteristics, this is unlikely: the low-contrast targets were less luminous than the high-contrast distractors, and no established early filtering mechanism selectively discards high-contrast stimuli. Thus, performance relied primarily on the participant’s ability to prevent extensive processing of distractors through a late filtering mechanism acting before the activation of the reading network and protecting verbal working memory. This is essential as verbal working memory must retain only the most recent words from the attended story to form coherent sentences rather than a meaningless mixture of unrelated words.

#### Additional Tasks Used for Functional Characterization

To further characterize the functional properties of specific iEEG sites, we also analyzed data collected during the following paradigms in the same participant cohort. **Visual Search Task (SEARCH)**. The task was adapted from the classic paradigm of Treisman and Gelade (***Treisman and Gelade, 1980***). Participants were asked to locate a target character embedded in an array of distractors. Each stimulus displayed a 6×6 grid containing 35 ‘L’ distractors and a single ‘T’ target in randomly assigned positions. Participants indicated the vertical position of the target (upper vs. lower half of the display) by pressing one of two buttons on a gamepad with the right index (upper) or middle (lower) finger. Two search conditions modulated task difficulty. In the EASY condition, the target was a grey ‘T’ among black ‘L’ distractors. In the HARD condition, all characters were grey. The blocked design consisted of 8 blocks of 12 stimuli (6 per condition) presented in pseudo-random order. Each stimulus appeared for 2500 ms, followed by a 1000 ms interstimulus interval. Stimuli were displayed centrally on a 19-inch computer screen positioned 60 cm from the participant. **Visual Oddball Task (VISU)**. In this task, participants pressed a button with their right index finger whenever a picture of a fruit appeared (20 targets total) (***Vidal et al., 2010***). Non-target stimuli consisted of images from eight categories: houses, faces, animals, landscapes, objects, pseudowords, consonant strings, and scrambled images (50 exemplars per category). All stimuli were matched for average luminance, except pseudowords and consonant strings, which were white letter strings on a black background. Images were presented within an oval aperture for 200 ms every 1000–1200 ms, in series of five images separated by 3-s pause periods during which participants could blink freely. Visual categories were pseudo-randomized across trials.

### Spectral and Statistical analysis

#### High-Frequency Activity extraction

High-Frequency Activity (HFA; 50–150 Hz) was used as a proxy for population-level spiking activity ( ***Buzsáki et al. (2012***),**?**) ). HFA was extracted from continuous bipolar iEEG signals following our standard procedure (***Perrone-Bertolotti and others, 2020***). Briefly: 1) Bandpass filtering — Signals were filtered into consecutive 10-Hz frequency bands (from 50–60 Hz to 140–150 Hz) using a fourth-order, two-way, zero-phase-lag Butter-worth filter. 2) Envelope extraction — The amplitude envelope of each filtered signal was computed using the Hilbert transform (***Quyen et al., 2001***) and downsampled to 64 Hz. 3) Normalization — Each envelope signal was normalized by dividing it by its mean amplitude over the entire recording session and multiplying the result by 100, allowing activity to be expressed as a percentage of the session mean. 4) Averaging across bands — The resulting normalized envelopes from all sequential bands were averaged to produce a single HFA time series, which by definition has a mean value of 100 across the recording session.

#### Statistical analysis of HFA relative to baseline and across conditions

To identify significant HFA modulations relative to the pre-stimulus baseline, we examined consecutive non-overlapping 25-ms windows spanning -300 ms to 800 ms relative to stimulus onset. For each window, we compared: a) the mean HFA within that window, b) to the mean HFA during the pre-stimulus baseline (-200 to 0 ms) using paired Wilcoxon signed-rank tests. Comparisons between experimental conditions (e.g., ATTEND vs. IGNORE words) were performed using a Kruskal–Wallis test applied independently to each 25-ms time window.

## Acknowledgments

This project has received funding from the European Union’s Horizon 2020 Research and Innovation Programme under Grant Agreement No. 720270 (HBP SGA1) and No. 785907 (HBP SGA2), and by EBRAINS 2.0 (HORIZON-INFRA-2022-SERV-B-01).

---

**Supplementary Figure 1.**
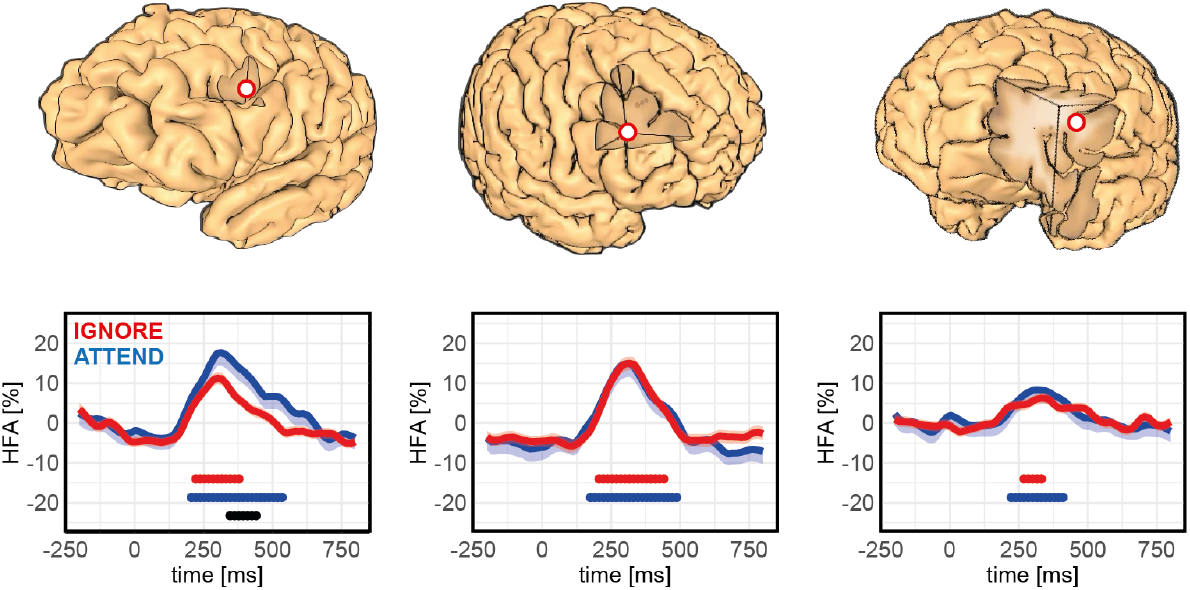
Early responses to attended and ignored words in the inferior frontal sulcus (IFS). Three additional IFS sites are shown on individual 3D brain reconstructions (red dots). All exhibited an Early Undifferentiated Response (EUR) during the Attentive Reading task. Colored horizontal bars indicate periods of significant high-frequency activity (HFA) increase relative to the pre-stimulus baseline (-200 to 0 ms), lasting more than 100 ms (Wilcoxon test, p < 0.001, uncorrected; stimulus onset at 0 ms). HFA is expressed as a percentage deviation from the mean HFA recorded through-out the experiment.

**Supplementary Figure 2.**
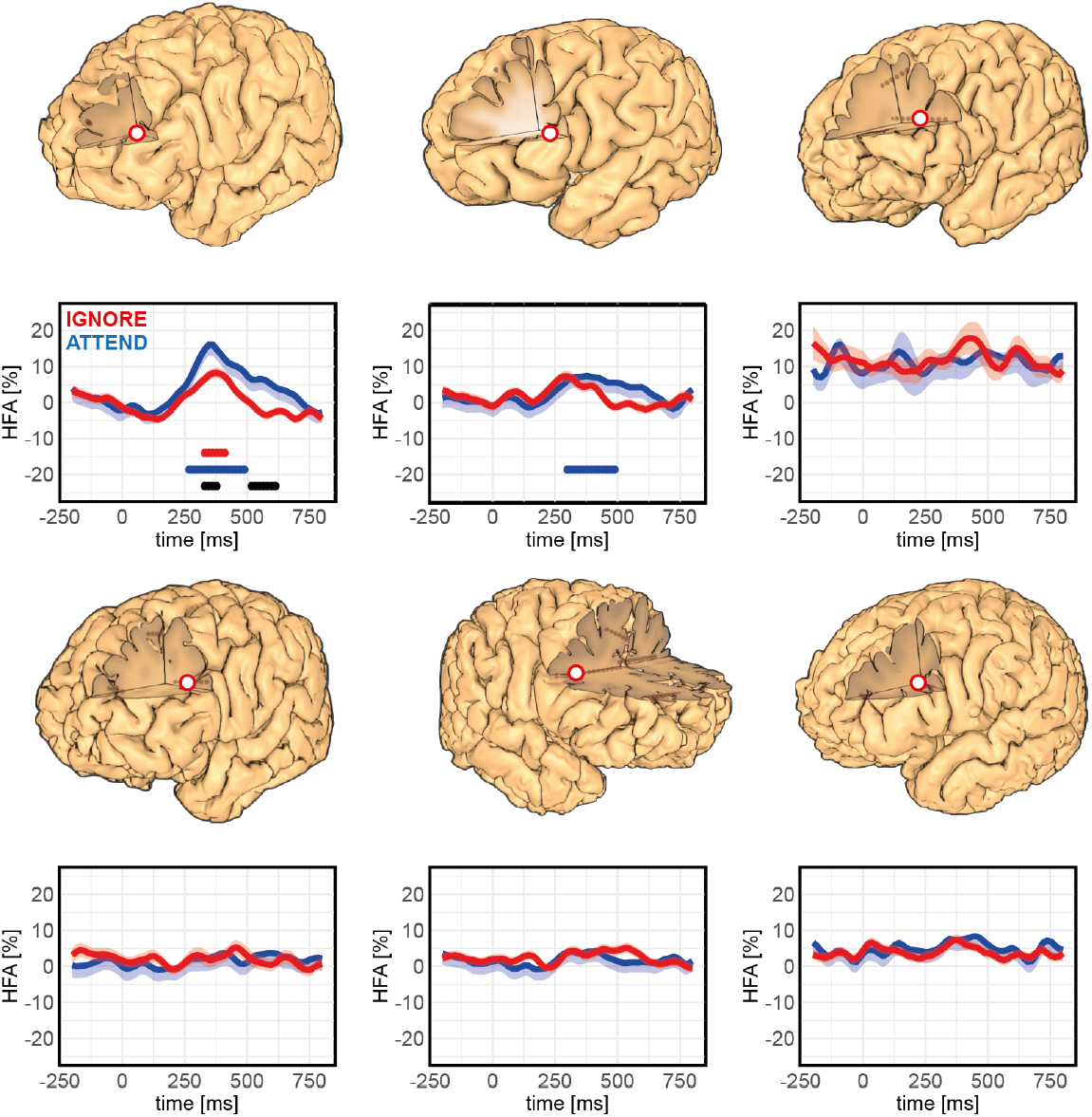
Non-responsive sites in the inferior frontal sulcus (IFS). Six IFS sites with weak or absent responses during the Attentive Reading task are shown on individual 3D brain reconstructions (red dots). No clear anatomical distinction could be established between these non-responsive sites and the responsive IFS sites shown in previous figures. Colored horizontal bars indicate periods of significant high-frequency activity (HFA) increase relative to the pre-stimulus base-line (-200 to 0 ms), lasting more than 100 ms (Wilcoxon test, p < 0.001, uncorrected; stimulus onset at 0 ms). HFA is expressed as a percentage deviation from the mean HFA recorded throughout the experiment.

**Supplementary Figure 3.**
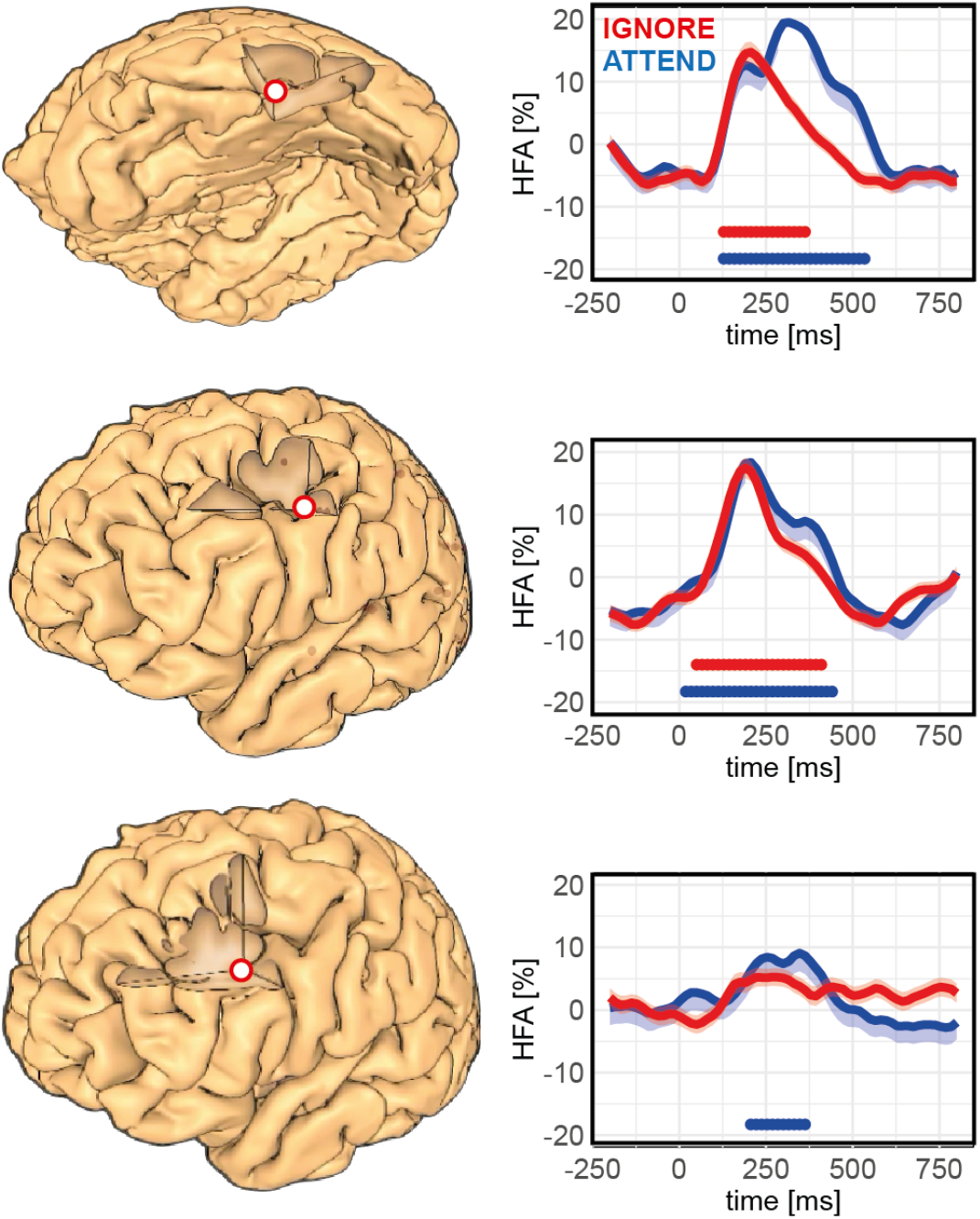
Comparative response dynamics in the VWFA, FEF, and IFS within a single participant. High-frequency activity (HFA) responses recorded during the Attentive Reading task are shown for three regions: the Visual Word Form Area (top), the Frontal Eye Field (middle), and the Inferior Frontal Sulcus (bottom). Precise electrode locations are displayed on the participant’s individual 3D brain model (red dots). Colored horizontal bars indicate periods of significant HFA increase relative to the pre-stimulus baseline (-200 to 0 ms), lasting more than 100 ms (Wilcoxon test, p < 0.001, uncorrected; stimulus onset at 0 ms). Black bars denote significant differences between attended and ignored conditions, also sustained for over 100 ms (Kruskal–Wallis test, p < 0.001, uncorrected). HFA is expressed as a percentage deviation from the mean HFA recorded throughout the experiment.

## References

Buzsáki G, Anastassiou CA, Koch C. The origin of extracellular fields and currents—EEG, ECoG, LFP and spikes. Nature Reviews Neuroscience. 2012; 13:407–420.

Chen L, Wassermann D, Abrams DA, Kochalka J, Gallardo-Diez G, Menon V. The visual word form area (VWFA) is part of both language and attention circuitry. Nature Communications. 2019; 10:5601. doi: 10.1038/s41467-019-13634-z.

Cole MW, Schneider W. The cognitive control network: Integrated cortical regions with dissociable functions. NeuroImage. 2007; 37:343–360.

Corbetta M, Patel G, Shulman GL. The reorienting system of the human brain: from environment to theory of mind. Neuron. 2008; 58:306–324. doi: 10.1016/j.neuron.2008.04.017.

Corbetta M, Shulman GL. Control of goal-directed and stimulus-driven attention in the brain. Nature Reviews Neuroscience. 2002; 3:201.

Deman P, others. IntrAnat Electrodes: A Free Database and Visualization Software for Intracranial Electroencephalographic Data Processed for Case and Group Studies. Frontiers in Neuroinformatics. 2018; 12.

Derrfuss J, Brass M, von Cramon DY. Cognitive control in the posterior frontolateral cortex: evidence from common activations in task coordination, interference control, and working memory. NeuroImage. 2004; 23:604–612.

Dosenbach NUF, Raichle ME, Gordon EM. The brain’s action-mode network. Nature Reviews Neuroscience. 2025; doi: 10.1038/s41583-024-00895-x.

Dosher BA, Lu ZL. Mechanisms of perceptual attention in precuing of location. Vision Research. 2000; 40:1269–1292. doi: 10.1016/S0042-6989(00)00019-5.

Duncan J. The demonstration of capacity limitation. Cognitive Psychology. 1980; 12:75–96.

Fiebelkorn IC, Kastner S. A Rhythmic Theory of Attention. Trends in Cognitive Sciences. 2019; 23:87–101. doi: 10.1016/j.tics.2018.11.009.

Huang YM, Yeh YY. Why does a red snake in the grass capture your attention? Emotion. 2011; 11:224–232. doi: 10.1037/a0022578.

Kastner S, Ungerleider LG. Mechanisms of visual attention in the human cortex. Annual Review of Neuroscience. 2000; 23:315–341. doi: 10.1146/annurev.neuro.23.1.315.

Lachaux JP, Rudrauf D, Kahane P. Intracranial EEG and human brain mapping. Journal of Physiology Paris. 2003; 97:613–628.

Mainy N, Jung J, Baciu M, Kahane P, Schoendorff B, Minotti L, Lachaux JP. Cortical dynamics of word recognition. Human Brain Mapping. 2008; 29:1215–1230.

Maunsell JHR, Treue S. Feature-based attention in visual cortex. Trends in Neurosciences. 2006; 29:317–322. doi: 10.1016/j.tins.2006.04.001.

Mercier MR, others. Advances in human intracranial electroencephalography research, guidelines and good practices. NeuroImage. 2022; 260:119438. doi: 10.1016/j.neuroimage.2022.119438.

Milham MP, others. The relative involvement of anterior cingulate and prefrontal cortex in attentional control depends on nature of conflict. Cognitive Brain Research. 2001; 12:467–473.

Nobre AC, Allison T, McCarthy G. Modulation of human extrastriate visual processing by selective attention to colours and words. Brain. 1998; 121:1357–1368. doi: 10.1093/brain/121.7.1357.

Ossandon T, others. Efficient ‘Pop-Out’ Visual Search Elicits Sustained Broadband Gamma Activity in the Dorsal Attention Network. Journal of Neuroscience. 2012; 32:3414–3421.

Peelen MV, Kastner S. Attention in the real world: toward understanding its neural basis. Trends in Cognitive Sciences. 2014; 18:242–250. doi: 10.1016/j.tics.2014.02.004.

Perrone-Bertolotti M, others. A real-time marker of object-based attention in the human brain: A possible component of a “gate-keeping mechanism” performing late attentional selection in the ventro-lateral prefrontal cortex. NeuroImage. 2020; 210:116574. doi: 10.1016/j.neuroimage.2020.116574.

Quyen MLV, Foucher J, Lachaux J, Rodriguez E, Lutz A, Martinerie J, Varela FJ. Comparison of Hilbert transform and wavelet methods for the analysis of neuronal synchrony. Journal of neuroscience methods. 2001 Oct; 111(2):83–98.

Ruland SH, Palomero-Gallagher N, Hoffstaedter F, Eickhoff SB, Mohlberg H, Amunts K. The inferior frontal sulcus: Cortical segregation, molecular architecture and function. Cortex. 2022; 153:235–256. doi: 10.1016/j.cortex.2022.03.019.

Sakai K. Task set and prefrontal cortex. Annu Rev Neurosci. 2008; 31(1):219–245. Publisher: Annual Reviews.

Seeley WW. The Salience Network: A Neural System for Perceiving and Responding to Homeostatic Demands. The Journal of Neuroscience. 2019; 39(50):9878–9882. doi: 10.1523/jneurosci.1138-17.2019.

Seeley WW, others. Dissociable Intrinsic Connectivity Networks for Salience Processing and Executive Control. Journal of Neuroscience. 2007; 27:2349–2356.

Talairach J, Tournoux P. Referentially oriented cerebral MRI anatomy: an atlas of stereotaxic anatomical correlations for gray and white matter. New York: Thieme; 1993.

Theeuwes J. Stimulus-driven capture and attentional set: Selective search for color and visual abrupt onsets. Journal of Experimental Psychology: Human Perception and Performance. 1994; 20:799–806.

Treisman A, Gelade G. A feature-integration theory of attention. Cognitive Psychology. 1980; 12:97–136.

Treue S, Martinez Trujillo JC. Feature-based attention influences motion processing gain in macaque visual cortex. Nature. 1999; 399:575–579. doi: 10.1038/21176.

Vecchio MD, Bontemps B, Lance F, Gannerie A, Sipp F, Albertini D, Cassani CM, Chatard B, Dupin M, Lachaux JP. Introducing HiBoP: A Unity-Based Visualization Software for Large iEEG Datasets. Journal of Neuroscience Methods. 2024; 409:110179. https://doi.org/10.1016/j.jneumeth.2024.110179, doi: 10.1016/j.jneumeth.2024.110179, publisher: Elsevier.

Vidal J, Ossandon T, Jerbi K, Dalal S, Minotti L, Ryvlin P, Kahane P, Lachaux JP. Category-specific visual responses: an intracranial study comparing gamma, beta, alpha, and ERP response selectivity. Frontiers in Human Neuroscience. 2010; 4:195.

Woolnough O, Forseth KJ, Rollo PS, Tandon N. Uncovering the functional anatomy of the human insula during speech. eLife. 2019; 8:e53086. doi: 10.7554/eLife.53086.

Zivony A, Eimer M. The diachronic account of attentional selectivity. Psychonomic Bulletin & Review. 2022; 29:1118–1142. doi: 10.3758/s13423-021-02023-7.

Zysset S, others. Color-word matching Stroop task: separating interference and response conflict. NeuroImage. 2001; 13:29–36.

